# Distribution of carbon monoxide (CO) in several tissues from Atlantic bottlenose dolphins (*Tursiops truncatus*)

**DOI:** 10.1101/2023.07.28.551019

**Authors:** Michael S. Tift, Kerryanne Litzenberg, Kayleigh M. Herrmann, Alicia T. Cotoia, Olivia N. Jackson, Tiffany F. Keenan, Kristi M. Kezar, Anna B. Pearson, William A. McLellan, D. Ann Pabst

## Abstract

Carbon monoxide (CO) is known as “The Silent Killer” due to its toxic effect at high concentrations, leading to an impairment in oxygen storage, delivery, and use. The cytotoxicity of CO is due to its high affinity for transition metals, such as iron, where CO outcompetes oxygen for the heme binding sites on hemoproteins in the body. CO is made *in vivo* in most organisms as a byproduct of heme degradation via heme oxygenase enzymes. Certain species of deep-diving marine mammals with high quantities of hemoproteins in blood and skeletal muscle have naturally elevated concentrations of CO in these tissues. To date, there exist few data on extravascular tissue CO content in wild animals. This study aims to characterize CO concentrations in nine different tissues from stranded Atlantic bottlenose dolphins (*Tursiops truncatus*). We found three tissues (liver, skeletal muscle, and spleen) have higher CO concentrations than other tissues. In a subset of samples from animals that tested positive for dolphin morbillivirus, the CO content in their kidney and liver was lower when compared to animals that tested negative. The mean CO concentration found in every tissue from dolphins was higher than those previously reported in healthy rodents. However, the skeletal muscle CO concentrations in dolphins from this study were much lower than those of deep-diving elephant seals. These results highlight the diversity and pattern of CO found in different tissues from bottlenose dolphins and continues to show that the heme oxygenase/carbon monoxide pathway appears to be critical for air-breathing divers.

## Introduction

It has been known for centuries that carbon monoxide (CO) is toxic to species that rely on hemoproteins to extract, store, and deliver oxygen (O_2_) to tissues for aerobic adenosine triphosphate (ATP) production (Douglas et al., 1912; Hopper et al., 2021). The toxicity arises from the high affinity CO has, compared to O_2_, for the heme binding site on several hemoproteins in the body. Most notably, CO has an affinity for heme binding sites on hemoglobin from adult humans that is approximately 218 times higher than O_2_ (Haab, 1990). If concentrations of CO rise in blood, the CO will outcompete O_2_ for the heme binding sites, resulting in a reduction in blood O_2_ carrying capacity that can lead to a reduction in mitochondrial O_2_ consumption, aerobic metabolism, and can eventually result in cell and tissue death if ATP demands are not met (Ernst and Zibrak, 1998).

Inhalation of CO from incomplete combustion of carbon-based fuels such as coal and gasoline are common exogenous sources that can result in large volumes of the gas entering the body, leading to toxic effects and one of the most common causes of poisoning in the United States (Raub et al., 2000). Surprisingly, the gas is also produced naturally in the body of most organisms as a byproduct of heme degradation (Wilks, 2002). Heme oxygenase enzymes (HO-1 and HO-2) catabolize heme, forming equimolar concentrations of biliverdin, with CO and free iron (Fe^2+^) as byproducts (Sjöstrand, 1949; Tenhunen et al., 1968). Therefore, every hemoglobin molecule that is fully degraded can result in the production of four molecules of CO being released into circulation. In humans and rodents, the highest quantity of CO in tissues other than blood has been found in the spleen and liver, the primary locations for erythrocyte and hemoglobin turnover (Vreman et al., 2005; Vreman et al., 2006). These two tissues are also known for having the highest heme oxygenase activity (Tenhunen et al., 1968). In conditions such as hemolytic anemia or sickle cell anemia, where patients are breaking down heme at a much faster rate, CO in the blood and exhaled breath of humans are elevated when compared to healthy individuals (Bensinger and Gillette, 1974; Coburn et al., 1966). In most cases, CO that enters the body through the lungs, or that is produced endogenously, will combine rapidly with hemoglobin in red blood cells, forming carboxyhemoglobin (COHb). However, CO also can interact with other hemoproteins (e.g., cytochrome-c-oxidase, soluble guanylyl cyclase) to impact cellular metabolism and pathways related to cell-signaling and cytoprotection (Cooper and Brown, 2008; Otterbein et al., 2000; Thom et al., 2000).

New research is shining light on the cytoprotective effects of heme oxygenase and carbon monoxide (Byrne et al., 2022; Hopper et al., 2021; Otterbein et al., 2016). Exposure to low or moderate concentrations of exogenous CO, or upregulation of endogenous CO production in the body via increased heme oxygenase activity, has resulted in cytoprotection in several tissues experiencing stress (Ryter, 2019). Specifically, several different types of diseases or injuries result in HO-1 to increase in quantity and activity in tissues, and this is often related to cytoprotective effects (Motterlini and Otterbein, 2010; Owens, 2010). Since heme is the primary source for endogenous CO production, it’s important to understand how variations in the quantity of heme stores in individuals relate to concentrations of the gas in the body. For example, certain species of marine mammals with naturally high levels of hemoproteins in blood (hemoglobin) and skeletal muscle (myoglobin) have concentrations of CO in their blood that resemble those seen in chronic cigarette smokers. Specifically, northern elephant seals (*Mirounga angustirostris*) and Weddell seals (*Leptonychotes Weddellii*) are two deep-diving phocids that have some of the highest reported hemoglobin concentrations and mass-specific blood volumes among mammals, with maximum COHb levels over 10% (Pugh, 1959; Tift et al., 2014). These findings are surprising given the dogma behind CO as only having negative impacts on O_2_ storage, transport, and use (Ernst and Zibrak, 1998). Therefore, the question remains, why do these animals that depend heavily on their ability to store large amounts of O_2_ for a diving lifestyle, also have high concentrations of a gas known to interfere with O_2_ storage and use? The upregulation of heme oxygenase/carbon monoxide pathway activity has been hypothesized to alleviate injuries associated with inflammation and apoptosis after exposure to hypoxia and/or ischemia reperfusion events; two common occurrences that marine mammal tissues routinely face as a side-effect of the physiological dive response (Tift et al., 2020). Rapid bradycardia and peripheral tissue ischemia at the onset of dives is followed by pronounced tachycardia and reperfusion to ischemic tissues once an animal surface, forcing many of their tissues to experience repeated ischemia-reperfusion events from diving, with no evidence of injuries (Allen and Vázquez-Medina, 2019). Therefore, marine mammals serve as an ideal model to understand mechanisms to avoid these injuries, where they may benefit from having modified heme oxygenase/carbon monoxide pathway to protect their cells and tissues (Tift and Ponganis, 2019).

There has also been evidence of the importance of the heme oxygenase/carbon monoxide pathway in non-diving species that experience and tolerate chronic hypoxia. For example, when comparing llamas (high-altitude adapted species) and sheep (domesticated lowland species), the activity of heme oxygenase proteins and production of CO was upregulated by pulmonary tissues of neonatal llamas that underwent gestation at high altitude, which resulted in an alleviation of pulmonary hypertension when compared to neonatal sheep that also underwent gestation at altitude (Herrera et al., 2008). It was specifically the enhanced CO, rather than nitric oxide (NO), that was hypothesized to provide protection against cell proliferation that contributes to pulmonary hypertension. This is similar to the cytoprotective effects of the heme oxygenase/carbon monoxide pathway seen in several other studies (Dubuis et al., 2005; Llanos et al., 2012; Zuckerbraun et al., 2006). When compared to mice, baseline *Hmox1* expression is also upregulated in the carotid body of naked mole rats, one of the most hypoxia tolerant mammals on the planet (Peng et al., 2022). However, despite high expression for the gene responsible for HO-1 production, the relative CO production rates in liver homogenates from naked mole rats did not change after exposure to one hour of hypoxia, while liver homogenates from mice experienced a drastic reduction in CO production rates (Peng et al., 2022). This is in contrast to an acute increase in COHb and HO-1 protein content in lungs of mice exposed to hypoxia (Carraway et al., 2000). Lastly, certain human populations living at high altitude in Tibet have several genes under positive selection that are suggested to be responsible for their ability to tolerate sustained hypoxia without increased risk of developing chronic mountain sickness from excessive erythrocytosis. One of the genes under positive selection in these populations is *HMOX2* (Simonson et al., 2010). It has been suggested that the modifications in this gene results in increased heme degradation activity by HO-2 in Tibetans, resulting in higher endogenous CO and reduced hemoglobin phenotype, protecting themselves from excessive erythrocytosis and other potential injuries associated with exposure to chronic hypoxia (Lin et al., 2023; Tift et al., 2020; Yang et al., 2016; Yu et al., 2022). Therefore, it appears that a modification in the heme oxygenase/carbon monoxide pathway is critical for species that are known to tolerate chronic hypoxia.

In order for CO to exert cytoprotection in tissues, the gas must dissociate from hemoglobin and interact with extravascular proteins and other molecules associated with cytoprotective pathways (R. Oliveira et al., 2016). Data on the quantity and diversity of extravascular CO in tissues (i.e., non-blood CO, COHb) is severely lacking. To date, extravascular tissue CO has been reported in tissues from healthy rodents and from deceased humans (Vreman et al., 2005; Vreman et al., 2006). Extravascular CO has also been recently reported in skeletal muscle from elephant seals which is over four times as high as that found in similar tissues from the healthy rodents and humans (Piotrowski et al., 2021). While most of the prior research has focused on measuring CO in blood, heme oxygenase proteins in tissues, and expression of heme oxygenase genes in relation to various conditions and treatments, very few studies have investigated the quantity and pattern of distribution of extravascular CO. Considering that CO must dissociate from hemoglobin to exert cytoprotective or signaling effects, it is important to understand how extravascular CO varies between species and with different conditions. In this study, we aimed to identify patterns in extravascular CO concentration between nine different organs from Atlantic bottlenose dolphins. We opportunistically sampled animals that stranded along the coast of North Carolina, USA between the years of 2006-2022. Considering CO production from heme oxygenase activity is known to increase from events of stress and disease, we also investigated whether animals that tested positive for dolphin morbillivirus (DMV) experienced differences in tissue CO concentrations.

## Methods

### Animals and sample collection

All samples were obtained from the University of North Carolina Wilmington Marine Mammal Stranding Program (NOAA SER Stranding Agreement to UNCW and IACUC #A1112-013, #A0809-019, #2006-015, #2003-013, #00-01). Tissues were collected from 27 adult common bottlenose dolphins, who were recovered stranded between the years of 2006 – 2022 on beaches throughout North Carolina, USA. Of these 27 individuals, 14 were male and 13 were female. Only animals with a Smithsonian Institution Code of 1 (live stranded and died naturally or by euthanasia; n=7) or 2 (fresh dead animal n=20) were used in this study (Geraci and Lounsbury, 1993). Necropsies were performed at UNCW on fresh carcasses or carcasses held in refrigerated storage overnight, and all tissues were collected and frozen within 1.5 days after initial contact with the animal. After sample collection of nine separate tissue types (brain, heart, kidney, liver, lung, skeletal muscle, spleen, testes, uterus), tissues were maintained in a - 80°C freezer to prevent freeze-thaw cycles and the premature release of CO prior to analysis. Twenty individuals in this study were also tested for infection with dolphin morbillivirus (DMV) through independent diagnostic labs (Marine Ecosystem Health Diagnostic & Surveillance Laboratory, One Health Institute, University of California Davis and the Zoological Pathology Program, University of Illinois at Urbana-Champaign (seven tested positive and thirteen tested negative).

### Quantification of CO in tissues

The extraction of CO and its quantification in tissues follows previously established methods (Byrne et al., 2022; Piotrowski et al., 2021; Vreman et al., 2005). Briefly, tissues were thawed and rinsed with chilled KH_2_PO_4_ buffer (pH 7.4) to remove any remaining excess blood and then placed into microcentrifuge tubes. Ice-cold de-ionized water was added to tissues samples at a (10% weight/weight ratio). With the tube on ice, the tissues were cut into smaller pieces using surgical scissors and thoroughly homogenized using an Ultra-Turrax T8 grinder and ultrasonic cell disruptor. Then, 10µL of the resulting tissue homogenate and 20 µL of 20% sulfosalicylic acid were added to purged amber chromatography vials to release CO into the headspace of the vial. Samples were stored on ice for at least 15 minutes before CO analysis. A CO-free carrier gas (Ultra-Zero Air; AirGas) was then used to push the headspace gas into the reducing compound photometer GC system (GC Reducing Compound Photometer, Peak Performer 1, Peak Laboratories LLC). Standard curves were generated daily for a certified calibration gas (1.02 ppm CO balanced with nitrogen: AirGas). Each tissue from each animal was analyzed in duplicate. The tissue CO concentrations were reported as pmol of CO per milligram wet weight tissue.

### Statistical Analyses

Data were analyzed using the statistical program JMP 16.0.0 (SAS Institute Inc.). The dataset containing the average CO concentration for each tissue was log transformed to meet normality assumptions for parametric tests. To examine the effects of DMV test results, sex, and Smithsonian Institution stranding code on the concentration of CO in tissues, we used two-sample t-tests. To examine the effect of sample storage duration on tissue CO concentrations, we used a linear regression model with stranding date as the independent variable and tissue CO concentration as the response variable. To test for significant differences in CO concentrations between tissues, we ran an ANOVA with tissue type as the independent variable, and subsequently performed a Tukey’s post-hoc analysis to determine significant differences between specific tissues. All mean values are reported as mean ± 1 standard deviation. Significance was determined at P<0.05.

## Results

Carbon monoxide concentrations were significantly different between the nine tissue types (F_8,60_ = 9.1; p < 0.0001). The highest mean CO concentrations were in liver (46.2 ± 18.3 pmol mg^-1^), skeletal muscle (45.4 ± 20.1 pmol mg^-1^), and the spleen (38.7 ± 14.4 pmol mg^-1^). The lowest mean CO concentrations were in uterus (10.7 ± 4.0 pmol mg^-1^), testes (11.4 ± 4.9 pmol mg^-1^), and the brain (12.2 ± 3.4 pmol mg^-1^) (Table 1; Fig. 1). For liver and kidney samples, tissue CO concentrations were significantly higher in animals that tested negative for DMV (51.3 ± 16.1 pmol mg^-1^ and 24.9 ± 7.9 pmol mg^-1^, respectively), compared to animals that tested positive for the virus (23.5 ± 3.4 pmol mg^-1^ and 14.4 ± 4.2 pmol mg^-1^, respectively) (Table 2; Fig. 2). There was no influence of sex or Smithsonian Stranding Code on tissue CO concentrations (Table 2; Fig. 3). There was a negative relationship between the average tissue CO content of all tissues with stranding date, but the relationship was weak due to low numbers of samples both early and late in the sampling period (p = 0.03, R^2^ = 0.07) (Fig. 4).

**Table 1:** Mean tissue carbon monoxide (CO) concentration in nine tissues from bottlenose dolphins. Values are representative of mean ± 1 Standard Deviation from 27 animals.

| Tissue Type | CO (pmol mg<sup>-1</sup>) |
| --- | --- |
| Brain | 12.2 ± 3.4 |
| Heart | 14.8 ± 7.8 |
| Kidney | 20.7 ± 8.4 |
| Liver | 46.2 ± 18.3 |
| Lung | 24.5 ± 18.1 |
| Skeletal Muscle | 45.4 ± 20.1 |
| Spleen | 38.7 ± 14.4 |
| Testes | 11.4 ± 4.9 |
| Uterus | 10.7 ± 4.0 |

**Table 2:** Statistical results of t-tests to investigate the impact of dolphin morbillivirus (DMV), sex, and Smithsonian Institution stranding code on extravascular tissue carbon monoxide (CO) in nine tissues from 27 bottlenose dolphins.

| Tissue | DMV result<br>(positive vs.<br>negative) | Sex | Stranding Code<br>(1 or 2) |
| --- | --- | --- | --- |
| Brain | P = 0.82 | P = 0.98 | P = 0.44 |
| Kidney | **P = 0.044** | P = 0.84 | P = 0.29 |
| Liver | **P = 0.012** | P = 0.18 | P = 0.06 |
| Lung | P = 0.43 | P = 0.95 | P = 0.76 |
| Skeletal Muscle | P = 0.15 | P = 0.89 | P = 0.36 |
| Spleen | P = 0.22 | P = 0.73 | P = 0.14 |
| Testes | P = 0.13 | N/A | P = 0.15 |
| Uterus | P = 0.90 | N/A | P = 0.23 |

**Figure 1:**
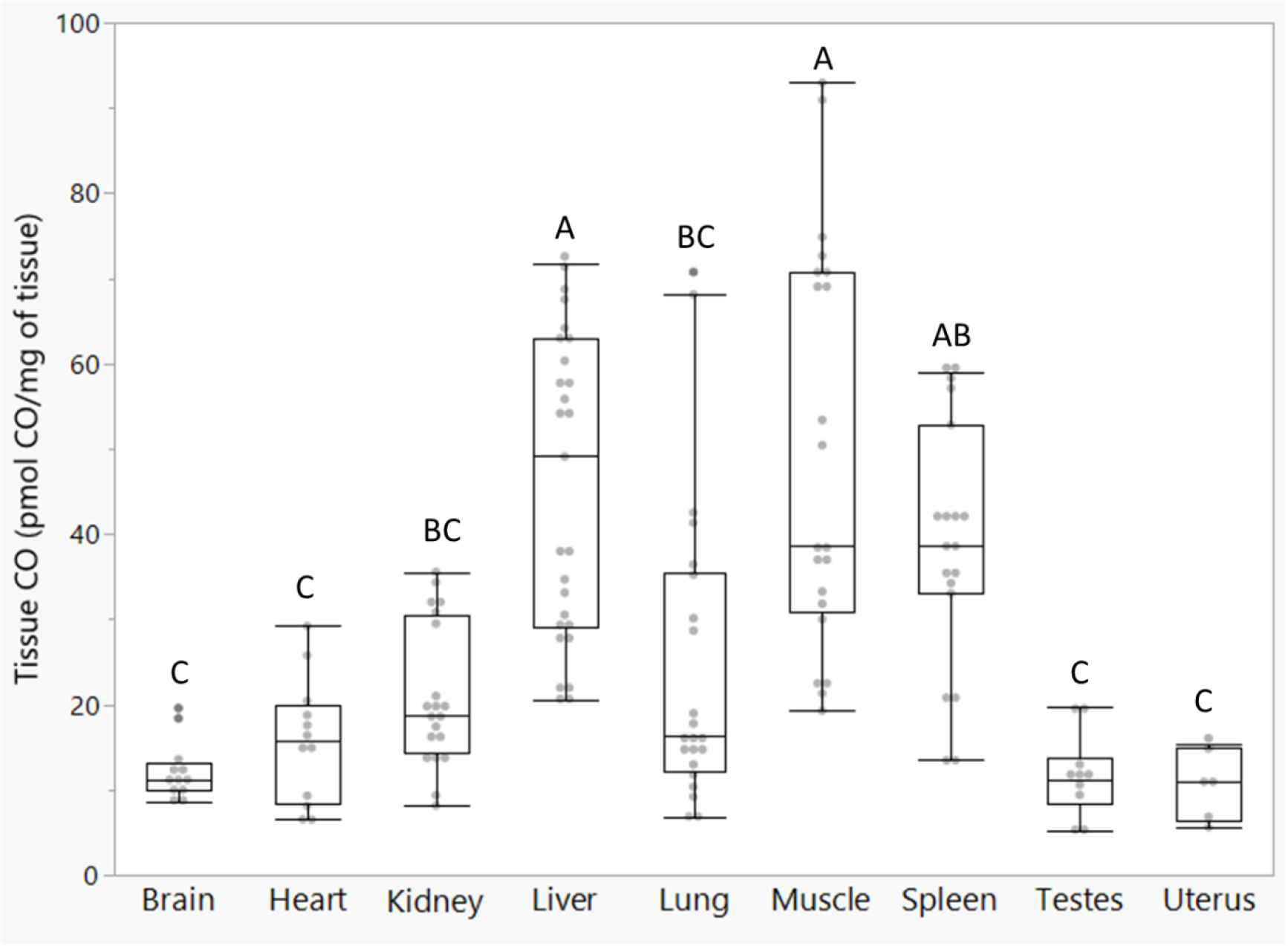
Tissue carbon monoxide (CO) concentration in extravascular tissues from bottlenose dolphins. Concentration of CO (pmol mg^-1^) from nine extravascular tissues from 27 animals. Boxes denote the upper (75%) and lower (25%) quantiles and the line in the box represents the median value for that tissue. Error bars represent the maximum and minimum values, excluding outliers (lower error bar = 1^st^ quartile – 1.5*[interquartile range]; upper error bar = 3^rd^ quartile + 1.5*[interquartile range]). Bars with different letters represent values that are significantly different from each other.

**Figure 2:**
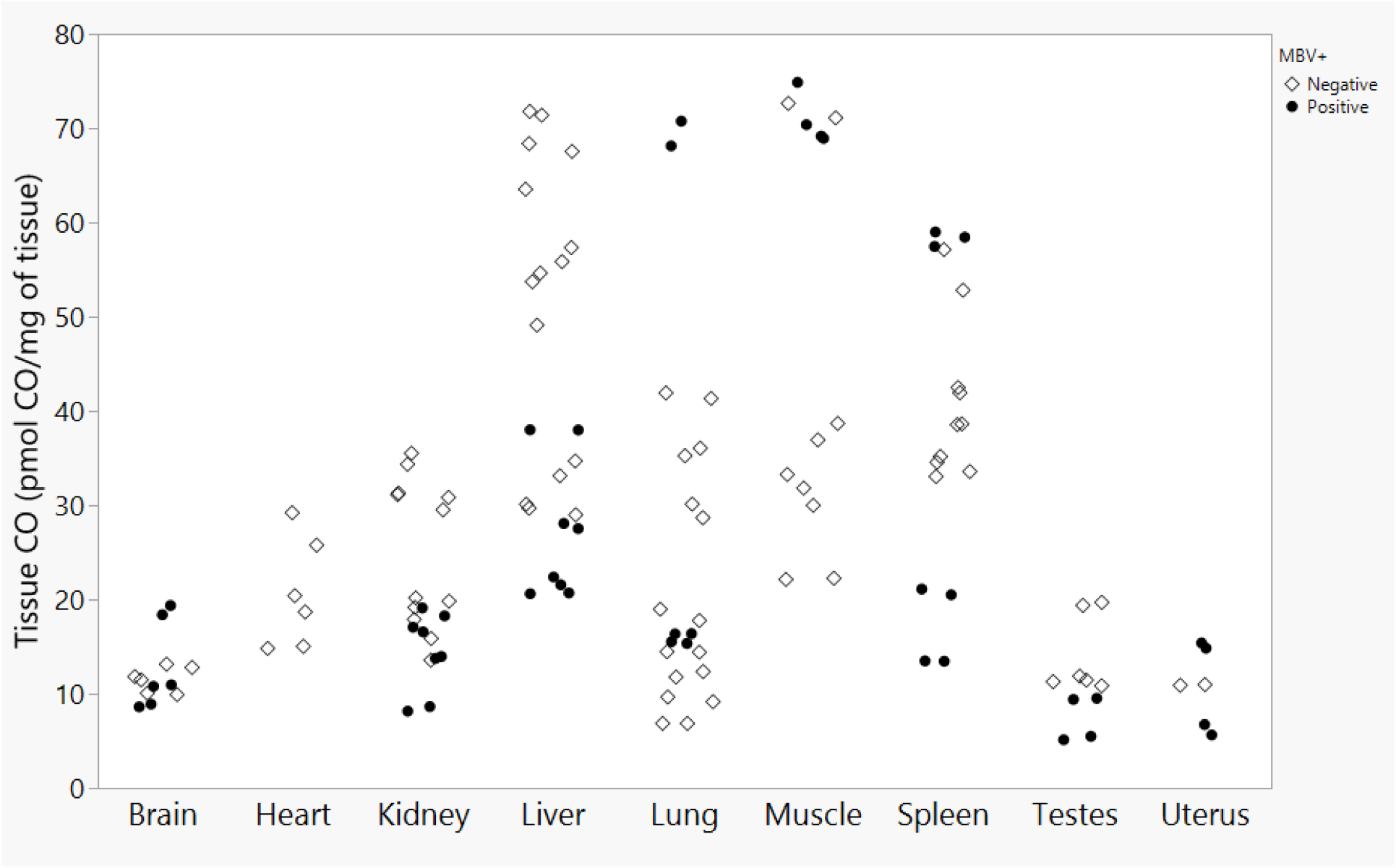
Tissue carbon monoxide (CO) concentration in extravascular tissues from bottlenose dolphins. Concentration of CO (pmol mg^-1^) from nine extravascular tissues in bottlenose dolphins, separated by disease status (negative = white diamonds; positive = black circles) for dolphin morbillivirus (DMV; MBV).

**Figure 3:**
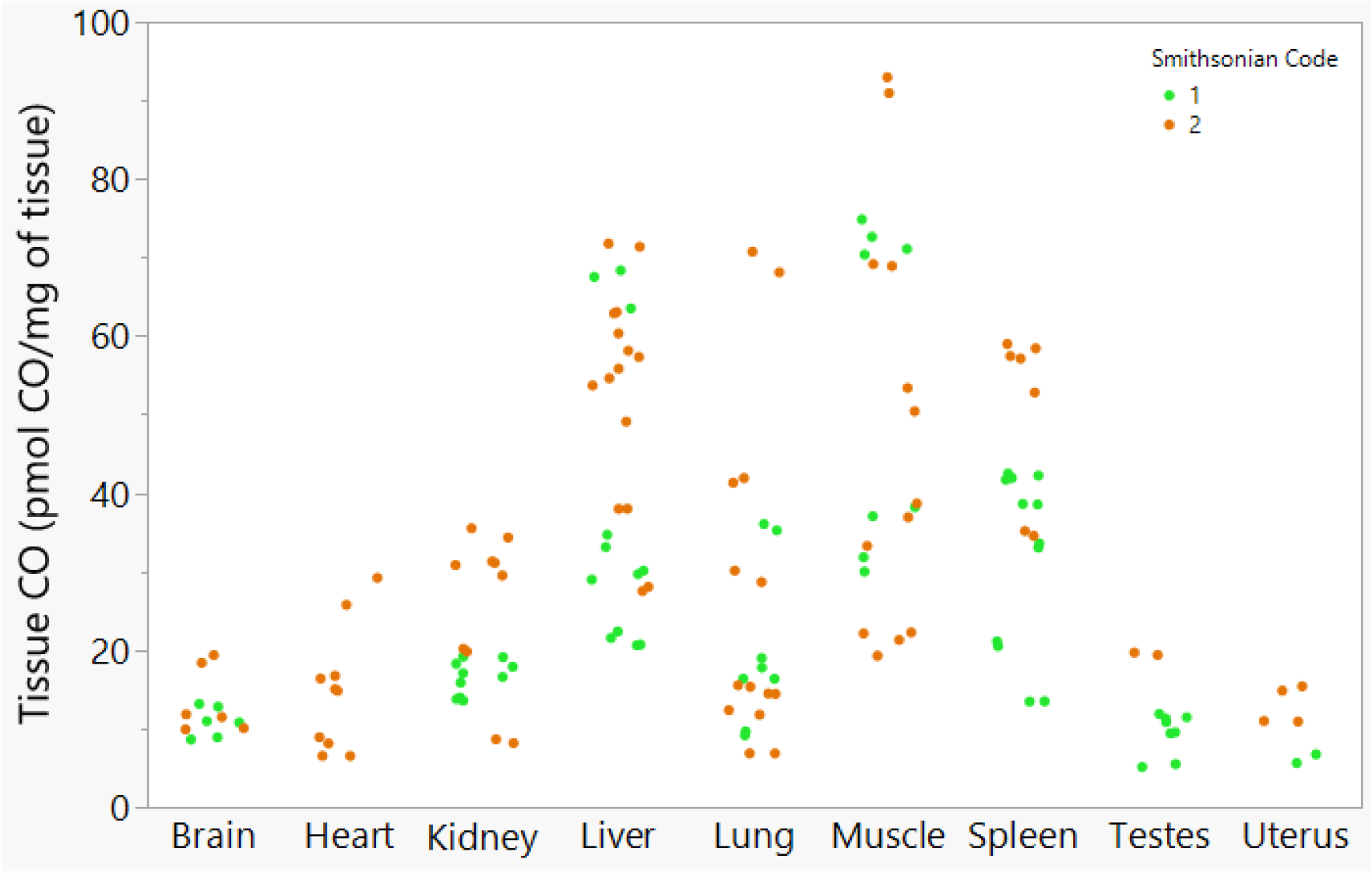
Tissue carbon monoxide (CO) concentration in extravascular tissues from bottlenose dolphins. Concentration of CO (pmol mg^-1^) from nine extravascular tissues in bottlenose dolphins. Animals are either assigned Smithsonian Code 1 (green; n = 7) or Code 2 (orange; n = 20).

**Figure 4:**
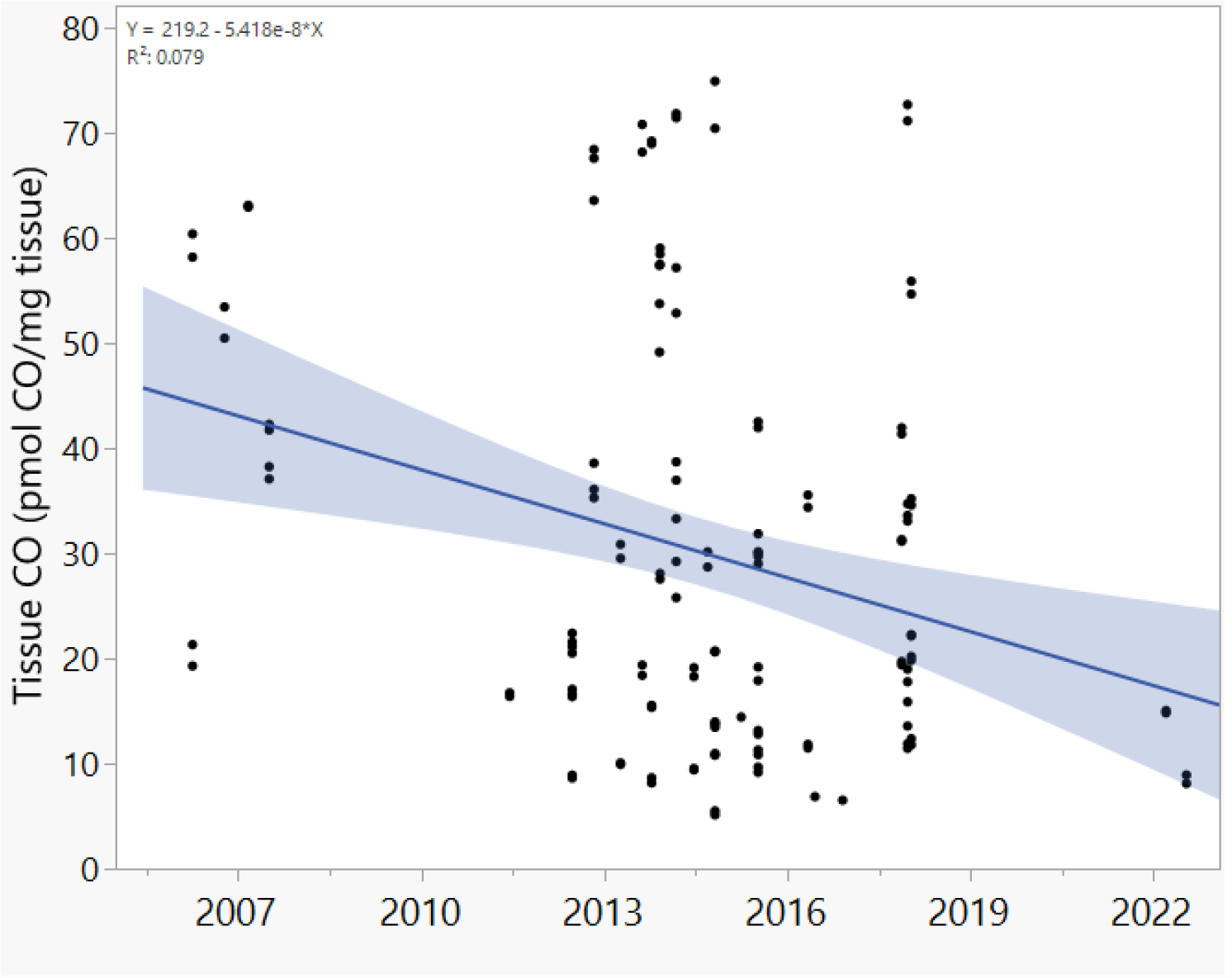
Tissue carbon monoxide (CO) concentrations in extravascular tissues from bottlenose dolphins collected from 2006 - 2022. Concentration of CO (pmol mg^-1^) from nine extravascular tissues in bottlenose dolphins. Each point represents the raw tissue CO value from each tissue, from each animal, plotted against the stranding date for that animal.

**Figure 5:**
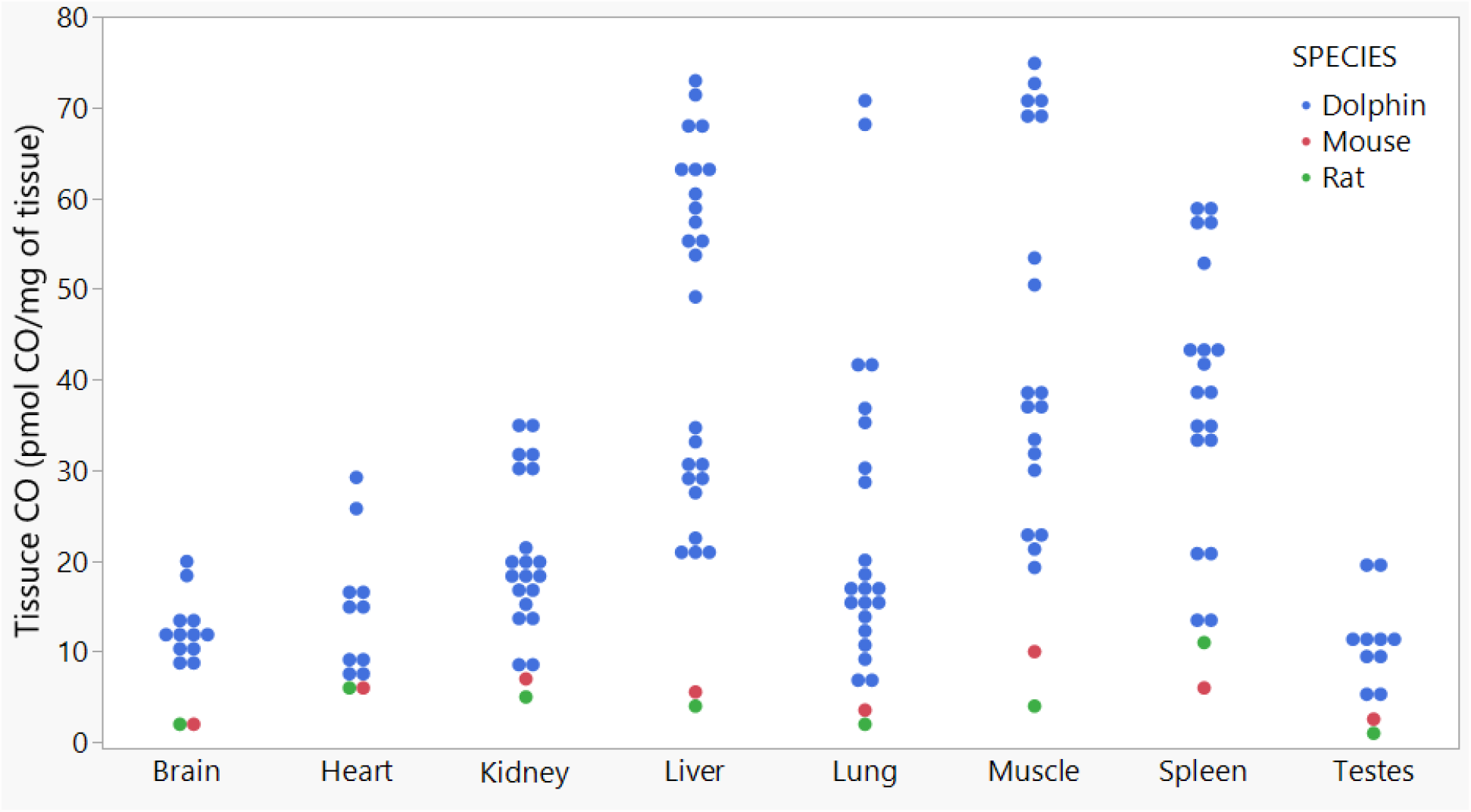
Tissue carbon monoxide (CO) concentration in extravascular tissues from bottlenose dolphins and healthy rodents. Concentration of CO (pmol mg^-1^) from eight extravascular tissues in bottlenose dolphins (blue), mice (red), and rats (green). Data for mice (n = 5) and rats (n = 3) are mean values from Vreman *et al*., 2005.

## Discussion

Similar to previous studies investigating extravascular CO concentrations, we show that the highest values were seen in liver, skeletal muscle, and spleen from bottlenose dolphins (Fig. 1) (Vreman et al., 2005; Vreman et al., 2006). The spleen and liver are known to be primary sites for red blood cell and hemoglobin degradation, and also have the highest levels of heme oxygenase activity (Vreman and Stevenson, 1988). Red blood cell turnover, and subsequent hemoglobin degradation, is known to be the source for ∼80% of endogenous CO production in healthy humans (Berk et al., 1974; Coburn et al., 1964; Tenhunen et al., 1968). Despite a much larger body mass than adult humans, bottlenose dolphins do have similar mass-specific blood volumes (∼70 mL kg^-1^), hemoglobin concentrations (∼13-15 g dL^-1^), and hematocrits (∼45%) (Dierauf and Gulland, 2001; Ridgway and Johnston, 1966). It is possible that the rate of turnover of red blood cells and hemoglobin in dolphins could be faster than in humans, resulting in greater endogenous CO production via heme oxygenase activity over time, resulting in higher tissue CO content. To date, the lifespan of red blood cells and hemoglobin turnover rates have never been reported in a marine mammal. Future studies should investigate blood CO concentrations in live animals and measure the contribution of erythrocyte lifespan and hemoglobin turnover rates towards endogenous CO production.

Other possible sources of endogenous CO production in the bottlenose dolphin could be the breakdown of hemoproteins other than hemoglobin via heme oxygenase enzymes, (e.g., myoglobin, cytochromes, catalases, peroxidases, soluble guanylate cyclase, and nitric oxide synthases) (Vreman et al., 2001). The half-life of myoglobin is not known for marine mammals, but has been reported to be 80-90 days in humans, which is shorter than their red blood cell lifespan (∼120 days) (Åkeson et al., 1960). The myoglobin content in skeletal muscle of marine mammals is known to be up to 30 times higher, when compared to their terrestrial counterparts (Ponganis, 2011; Ponganis, 2015). The myoglobin content in skeletal muscle of dolphins is approximately 7 times higher than that of adult humans (Kielhorn et al., 2013; Möller and Sylven, 1981). This high concentration of myoglobin serves as an O_2_ reservoir to increase total body O_2_ storage capacity in diving animals but can also act as a source and/or sink for endogenous CO via heme oxygenase enzymes. The percent of total body weight made of muscle is ∼2.5 times higher in bottlenose dolphins than in humans, and they have roughly 8-10 times more myoglobin in primary locomotory muscles (Goforth, 1987; Mallette et al., 2016; Noren and Williams, 2000; Velten et al., 2013). When compared to healthy mice, rats, and humans, skeletal muscle from bottlenose dolphins had 4.5, 10, and 3X higher CO concentrations, respectively (Vreman et al., 2005; Vreman et al., 2006). This could be due to dolphins having much higher myoglobin concentration in their skeletal muscle, or it could represent higher amounts of heme oxygenase activity in the skeletal muscle. A comparison of CO content in skeletal muscle from other species with different myoglobin concentrations, together with heme oxygenase activity measurements, and quantification of heme oxygenase protein content, would help to address this question.

The kidney and liver from dolphins that tested positive for DMV had significantly lower concentrations of CO than the same tissues from animals that tested negative (Fig. 2). This result was surprising, as it has been well documented that HO-1 is a stress-inducible enzyme that is upregulated in quantity and activity in many animals and humans that experience stress and/or disease (Owens, 2010). Cetaceans that have contracted DMV can often experience encephalitis and bronchointerstitial pneumonia, which could trigger an immune response in the brain and lung (Di Guardo and Mazzariol, 2016). Considering the importance of protein structure in the function of enzymes, it is essential to remember that the amino acid structure of the HO-1 enzyme was found to be conserved between humans and bottlenose dolphins, with >82% similarity (Reyes-Ramos et al., 2021). Despite this structural resemblance, leukocytes from bottlenose dolphins exposed to a proinflammatory challenge had significantly higher HO-1 activity and expression of *Hmox1* compared to human leukocytes (Reyes-Ramos et al., 2023). There were no distinct differences in CO content of the brain and lungs of animals that tested negative or positive for DMV from this study. It is possible that the reduced quantity of CO seen in the kidney and liver from animals that tested positive for DMV represent a reduction in heme oxygenase activity and associated CO production in those tissues. This matches results from previous studies that have shown an increase in heme oxygenase/carbon monoxide pathway activity at the early onset of certain pathologies, followed by a reduction in the pathway activity once the disease transitioned into a chronic response (Carraway et al., 2002; Cronje et al., 2004). It is not possible to determine when the animals from this study contracted DMV. Therefore, future studies should investigate differences in the heme oxygenase/carbon monoxide pathway of specific tissues and in the face of different stressors and in relation to the duration and severity of stressor exposure.

In a resting state (i.e., baseline), the percent of COHb stays constant in mice, rats, and humans, which suggests that the animals are producing and removing equal proportions of CO, preventing CO from rapidly changing. While tissue CO concentration was highest in the liver, skeletal muscle, and spleen of the bottlenose dolphin, it is worth noting that every tissue used in this study had higher mean CO concentrations than those previously reported in healthy rodents (Fig. 4) (Vreman et al., 2005). This could be due to differences in mass-specific rates in metabolism, CO production, and/or CO removal rates. For example, mice have a significantly reduced red blood cell lifespan (∼55 days), when compared to humans (∼120 days), resulting in a mass-specific CO production rate that is ∼5X higher than that of adult humans (Coburn, 2012; Vreman and Stevenson, 1988). Despite this difference in the magnitude of mass-specific CO production rates, the smaller mammals might also be more effective at removing CO from their body to prevent the gas from accumulating in blood and tissues (Vreman et al., 2005). Supporting this hypothesis is the fact that the smaller mammals have much higher heart rates and respiratory rates to meet their higher mass-specific metabolic demands, which could help with the removal of endogenous CO via the lungs (West, 1974).

Another important factor to consider that could help to explain higher amounts of CO in tissues of bottlenose dolphins is the impact of breath-holding on the removal of CO from the body. Their average dive duration is around 1 minute (max = 7.5 minutes), and depth is 20 m (max =390 m), spending between 82 – 93% of their lives submerged (Klatsky et al., 2007; Mate et al., 1995; Ridgway, 1986). Considering that CO is primarily removed from the body through the lungs, chronic breath-holding behaviors could result in a buildup of endogenous CO over time (Henderson and Haggard, 1920). Higher CO content in blood and exhaled breath has been seen in human patients that experience sleep apnea, and levels of CO have been positively related to the apnea-hypopnea index and percentage of total sleep time that patients experienced hypoxemia (Azuma et al., 2017; Kobayashi et al., 2008). It was presumed that this increase in endogenous CO in humans is attributable to the stresses associated with sleep apnea (e.g., hypoxemia, blood pressure variations, oxidative stress), rather than breath holding itself, that led to an increase in the stress-inducible heme oxygenase-1 activity and CO production. However, the exact mechanism behind elevated CO in patients with sleep apnea or in animals that experience repeated breath-holds, has yet to be determined. Some marine mammals are known to hyperventilate during surface intervals that follow dives, and dolphins are also known to be able to exchange 75-90% of their total lung volume within a fraction of a second (Andrews et al., 2000; Irving et al., 1941; Kooyman and Cornell, 1981; Piscitelli et al., 2010; Ridgway et al., 1969). Together, these breathing patterns and unique anatomy provide mechanisms for the exchange of large volumes of gases in marine mammals, which should be further investigated in relation to the removal rate of CO.

This study highlights yet another marine mammal species with high quantities of endogenous CO (Piotrowski et al., 2021; Pugh, 1959; Tift et al., 2014). Extravascular CO content was much higher in the tissues from bottlenose dolphins, when compared to healthy rodents (Vreman et al., 2005), and also when compared to certain tissues in humans (Vreman et al., 2006). The source of this CO is currently unknown, but it is hypothesized to be a byproduct of the catabolism of heme, by heme oxygenase enzymes. The naturally occurring high concentration of hemoproteins in the blood (hemoglobin) and muscle (myoglobin) of marine mammals provides a large quantity of heme for CO production. Future studies should investigate the turnover rates of specific hemoproteins in these species to understand their contribution to endogenous CO production. It is possible that the maintenance of high concentrations of extravascular CO is advantageous for diving animals, and other animals exposed to repeated events of hypoxia and/or ischemia reperfusion events, given the cytoprotective properties that low-to-moderate amounts of CO have shown for certain tissues (Motterlini and Otterbein, 2010). The reduced quantity of extravascular CO in the liver and kidney of animals with DMV should be further investigated, as CO is typically upregulated in other animals with disease. COHb in blood of animals that also have extravascular CO quantified could also provide evidence on the distribution of CO throughout the body.

## Acknowledgements

We thank Karen Clark (NC Wildlife Resources Commission); Paul Doshkov, Will Thompson and colleagues (National Park Service); Marina Piscitelli-Doshkov (NC Aquarium at Roanoke Island); Bruce Ferrier (NOAA); and Logan Arthur, Ryan McAlarney, Erin Cummings, Emily Singleton, and additional members of the UNCW Marine Mammal Stranding Program for their collegial assistance in stranding investigations. Work was carried out under a NOAA Stranding Agreement to UNCW and under UNCW IACUC No. 2003-013, 2006-015, A0809-019, and A2021-013 protocols.

## Funding

This work was supported by NSF grant #1927616, and NOAA Prescott grants (NA20NMF4390119 and NA21NMF4390398).

## Notes

### Competing Interest Statement

The authors have declared no competing interest.

